# Modulation of the *DA1* pathway in maize shows that translatability of information from Arabidopsis to crops is complex

**DOI:** 10.1101/2022.03.22.485335

**Authors:** Pan Gong, Kirin Demuynck, Jolien De Block, Stijn Aesaert, Griet Coussens, Laurens Pauwels, Dirk Inzé, Hilde Nelissen

## Abstract

Modern agriculture is struggling to meet the increasing food, silage and raw material demands due to the rapid growth of population and climate change. In Arabidopsis, DA1 and DAR1 are proteases that negatively regulate cell proliferation and control organ size. DA1 and DAR1 are activated by ubiquitination catalyzed by the E3 ligase BIG BROTHER (BB). Here, we characterized the *DA1, DAR 1* and *BB* gene families in maize and analyzed whether perturbation of these genes regulates organ size similar to what was observed in Arabidopsis. We generated *da1_dar1a_dar1b* triple CRISPR maize mutants and *bb1_bb2* double mutants. Detailed phenotypic analysis showed that the size of leaf, stem, cob, and seed was not consistently enlarged in these mutants. Also overexpression of a dominant-negative *DA1^R333K^* allele, resembling the *da1-1* allele of Arabidopsis which has larger leaves and seeds, did not alter the maize phenotype. The mild negative effects on plant height of the *DA1^R333K^_bb1_bb2* mutant indicate that the genes in the DA1 pathway may control organ size in maize, albeit less obvious than in Arabidopsis.

## 1. Introduction

The increasing global food demand caused by a rapidly growing population is the major issue facing modern sustainable agriculture and improving crop biomass and seed yield is the key to solve this problem. One of the worldwide grown crops, maize (*Zea mays*), is amongst the most important sources of food, silage, and bioenergy. In recent decades, genetic engineering techniques have been demonstrated to be powerful means to improve plant organ size in various crops, such as rice, wheat, and maize [1]. Thus, deciphering genes and networks that regulate organ size in maize may significantly contribute to food security.

Plant organ size is mainly determined by the combination of cell proliferation and cell expansion which contribute to the total cell number and the final cell size, respectively [2]. Cell proliferation determines early organ growth while all cells are relatively constant and have small sizes [3]. Then, cells gradually exit the mitotic cell cycle and transit to differentiation and expansion [4, 5]. Cell proliferation and cell expansion are regulated by transcriptional regulators [6], phytohormones [7] and post-transcriptional modifications [8]. At the post-transcriptional level, ubiquitination plays a prominent role in controlling organ size. Three types of enzymes participate in this catalytic process: the ubiquitin-activating enzyme (E1), the ubiquitin-conjugating enzyme (E2), and the ubiquitin ligase (E3), which facilitates the transfer of ubiquitin from the E2-ubiquitin intermediate to the substrate protein [9].

The ubiquitin-dependent protease DA1 (DA means “large” in Chinese) was first identified in Arabidopsis in a genetic screen for mutants with larger seeds [10]. DA1 harbors two ubiquitin interacting motifs (UIM) [11], a LIM domain which is essential for its protease activity and a conserved C-terminal region which contains the DA1 peptidase motif [12]. The dominant-negative Arabidopsis mutant *da1-1* (*DA1^R358K^*) contains a point mutation in the C-terminal region converting the 358^th^ arginine into lysine. The *da1-1* showed enlarged leaves, petals, and seeds due to a prolonged cell proliferation duration [10, 12, 13]. Overexpression of the *DA1^R358K^* dominant-negative allele also mimicked the *da1-1* phenotype [10]. Furthermore, overexpression of Arabidopsis *DA1^R358K^* in *Brassica napus* caused larger organs, including seeds, cotyledons, leaves, flowers, and siliques [15]. Down-regulating the wheat *DA1* homologue, *TaDA1*, also showed an increase in kernel size and weight, while overexpression of *TaDA1* resulted in the opposite phenotypes [16]. Overexpression of the soybean *DA1* did not alter the seed size but increased salt tolerance [17]. Moreover, overexpression of the dominant-negative allele of *DA1* in maize inbred line DH4866 did not promote final leaf size but appeared to increase seed size [18]. In Arabidopsis, the expression of *DA1* was increased by abscisic acid, indicating that the *DA1* may participate in response to abiotic stresses [10].

BIG BROTHER/ENHANCER 1 OF DA1 (BB/EOD1, referred to from hereon as BB) is a RING E3 ligase that interacts with DA1 and negatively regulates cell proliferation [19, 20]. The RING domain of E3 ubiquitin ligases binds to the E2 and is essential for the catalytic activity [21]. The Arabidopsis *bb* mutants showed increased organ size due to a prolonged cell proliferation period while overexpression of *BB* reduced organ growth [13, 20]. Moreover, the double *da1-1_bb* mutant had synergistic effects on promoting organ size through a substantial increase in cell number [10, 12, 13]. The RING E3 ligase DA2 also physically interacts with DA1 [19]. Loss-of-function *da2* mutants produced larger organs such as leaves, flowers and seeds by affecting cell proliferation [19]. DA1 and DA2 had a synergistic effect on organ growth and the double mutant *da1-1_da2-1* generates larger seeds and petals than WT and the single mutants [19]. The *bb_da2* double mutants showed an additive effect on organ size, suggesting that these two E3 ligases worked independently [10, 12, 13]. DA1 cleaves various target proteins when it is ubiquitinated through BB or DA2. As a negative feedback loop, DA1 also cleaves these E3 ligases [12]. At present, several DA1 substrates were found in Arabidopsis, such as the UBIQUITIN-SPECIFIC PROTEASE 15 (UBP15) [22] and the plant-specific TCP (named after the first 4 characterized members, namely TEOSINTE BRANCHED1 (TB1) from maize, CYCLOIDEA (CYC) from snapdragon (Antirrhinum majus), as well as PROLIFERATING CELL NUCLEAR ANTIGEN FACTOR1 (PCF1) and PCF2 from rice (Oryza sativa)) transcription factors TCP14 and TCP15 [12]. In Arabidopsis, disruption of *UBP15* resulted in smaller plants with narrow and serrated leaves, smaller flowers, and shorter siliques [23]. Conversely, overexpression of *UBP15* increased organ size by promoting cell proliferation [22, 23]. In rice, overexpression of the rice *OsUBP15* slightly increased the grain size [24]. TCP14 and TCP15 interact with DA1, DAR1 and DAR2 and act redundantly in regulating organ growth [14]. The *tcp14_tcp15* double mutants showed decreased inflorescence height, pedicel length and internode length [25]. The growth defects of triple *da1ko1_dar1-1_dar2-1* mutants were partially restored in the pentuple *tcp14-3_tcp15-3_da1-ko1_dar1-1_dar2-1* mutant [14].

There are seven *DA1-related* genes (*DARs*) in Arabidopsis and *DAR1* and *DAR2* genes are the most closely related family members of *DA1*. The loss-of-function T-DNA allele of *DA1* (*da1-ko1*) did not cause an obvious growth phenotype, nor did T-DNA insertion alleles in *DAR1* (*dar1-1*) or *DAR2* (*dar2-2*). However, the double *da1-ko1_dar1-1* knockout mutants showed increased organ size similar to the phenotype of *da1-1* [10]. The triple *da1-ko1_dar1-1_dar2-2* mutants have a decreased leaf size but enlarged petal and seed size. Overexpression of either *DA1, DAR1* or *DAR2* in the *da1-ko1_dar1-1_dar2-2* triple mutant restores its phenotype back to WT, suggesting that *DA1, DAR1* or *DAR2* are functionally redundant [14].

Here, we tested the involvement of the maize *DA1* pathway under well-watered and drought conditions. We overexpressed a similar dominant-negative mutation of the maize *DA1^R333K^* in the inbred line B104 which did not result in enhanced organ size. Next, we generated the *da1_dar1a_dar1b* triple knockout and *bb1_bb2* double knockout mutants by using CRISPR/Cas9 genome editing. Unfortunately, the leaf length, plant height and seed size did not show consistent significant changes compared with WT. Furthermore, the cross between *DA1^R333K^* and the *bb1_bb2* and the *DA1^R333K^_bb1_bb2* mutant showed significantly decreased plant height, suggesting that the *DA1* and *BB* genes affect organ size in maize.

## 2. Material and methods

### 2.1 Growth conditions in growth chamber and greenhouse

Plants for leaf growth monitoring were grown under growth chamber conditions with controlled relative humidity (55%), temperature (24 °C day/18°C night), and light intensity (170–200 μmol/m^2^/s photosynthetic active radiation at plant level) provided by a combination of high-pressure sodium vapor (RNP-T/LR/400W/S/230/E40; Radium) and metal halide lamps with quartz burners (HRI-BT/400W/D230/E40; Radium) in a 16-h/8-h (day/night) cycle. Plants for adult plant trait characterization were grown under controlled greenhouse conditions (26 °C/22 °C, 55% relative humidity, the light intensity of 180 μmol/m^2^/s photosynthetic active radiation, in a 16-h/8-h day/night cycle).

### 2.2 Analysis of protein sequence

The PLAZA protein database was searched with the BLASTP method using the amino acid sequences of the Arabidopsis DA1 (AT1G19270) and BB (AT3G63530). Proteins of DA1 families and BB in Arabidopsis, rice and maize were used to generate phylogenetic trees. All alignments of protein sequences were generated with ClustalW and phylogenetic trees were generated using maximum likelihood with applying 100 bootstraps method of MEGA X.

### 2.3 Constructs generation

The coding sequences of *DA1* (GRMZM2G017845) (containing a 998^th^ guanine replaced by adenine) flanked by AttB1/AttB2 Gateway recombination sites was synthesized and ligated into pDONR221 entry vectors using Gateway BP reactions. The *DA1* entry vector together with the UBIQUITIN (pUBIL) promoter [26] were ligated into the pBb7m24GW destination vector [27] to generate the pUBIL: *DA1^R333K^* (*DA1^R333K^*) expression vector by Multisite Gateway recombination (Invitrogen). This vector is codon-optimized for monocots and contains a Bar selection marker for the selection of transgenic plants using phosphinotricin [27]. CRISPOR [28] was used to determine possible gRNAs and their off-target in the coding sequence of *DA1, DAR1* and *BB* genes. Primers were designed that matched with the gRNA sequences (Table S2), after which PCR was performed on the pCBC-MT1T2 plasmid, resulting in a fragment containing the desired target sites and the correct sites for ligation into the pBUN411-Sp destination vector [29, 30] using Golden Gate cloning (Invitrogen). The ligation sites from different vectors were verified by Sanger sequencing.

### 2.4 Maize transformation and genotyping

Immature embryos of the maize inbred line B104 were transformed by *Agrobacterium tumefaciens* cocultivation [26]. In short, immature B104 embryos were co-cultivated with *A. tumefaciens* (EHA101) for 3 days followed by 1-week growth on non-selective medium. Transformed embryogenic calli were subsequently selected on increasing concentrations of phosphinotricin. After shoot induction from the selected calli, transgenic T_0_ shoots were transferred to soil. At maturity, these T_0_ shoots were backcrossed with B104 WT, resulting in a collection of BC_1_ seeds from multiple independent transgenic events. For the overexpression lines, the segregating population of 1:1 non-transgenic: transgenic plants were used for phenotyping and biomass generation. For the genome editing lines, the offspring containing the heterozygous mutant alleles that did not contain the T-DNA harboring CRISPR/Cas9 were self-crossed to identify WT plants and homozygous mutants. Then, the segregating population was self-pollinated for phenotypic analysis in the next generation. Screening for mutations and genotyping of CRISPR mutants was performed by sequencing the genomic region containing both target sites of each gene after PCR amplification. The primers used for PCR were listed in Table S2.

### 2.5 Quantitative RT-PCR

Expression analysis of *DA1* was checked by qRT-PCR. Total RNA was extracted from each repeat using Trizol (Life Technologies, Invitrogen) and DNA was removed by RQ1 DNase (Promega) treatment. Preparation of cDNA was performed using the iScript cDNA synthesis kit (BioRad) according to the manufacturer’s recommendations starting with 1μg of RNA. qRT-PCR was performed with the LightCycler 480 Real-Time SYBR Green PCR System (Roche), used primers are listed in Table S2. For qPCR on maize samples, we used *18S RNA* as the housekeeping gene.

### 2.6 Measurements of maize agronomy traits

The plant height was measured from soil base to highest leaf collar at silking stage. Ear leaf is the leaf that covers the main ear. Ear leaf is the leaf that covers the main ear. Ear leaf length was measured from the leaf auricle to the leaf tip, and the middle part of the blade determined its width. Stem width was measured at the internode base where the main ear grew. Stem width1 and 2 represent two diameters of the elliptic internode. 30 seeds in the middle of each cob were placed flat on paper, and pictures were taken. The average seed area was analyzed by using ImageJ.

### 2.7 Drought treatments

For mild drought treatment, the water content was allowed to drop to a soil water content of 70% of the well-watered conditions, corresponding to a soil water potential of −0.518 and −0.023 MPa, respectively. For drought treatment, when leaf 3 appears, no water was applied to the shoots under drought conditions.

## 3. Results

### 3.1 Identification of *DA1, DAR1* and *BB* genes in maize

The *DA1* gene family in maize has six members and all of them contain the LIM domain and the conserved C-terminal region [18]. GRMZM2G017845 was the closest homologue to the Arabidopsis *DA1* and was identified as the maize *DA1* [18]. The phylogenic tree of *DA1* families showed that two other maize genes clustered with the *AtDAR1*, and we defined the GRMZM2G099328 as *DAR1a* and the GRMZM2G342105 as *DAR1b*. Moreover, two maize *DAR2* homologues (GRMZM2G160198 and GRMZM2G151934) were found in the *AtDAR2* clade (Fig. S1A) but these are distantly related to the maize *DA1*, *DAR1a* and *DAR1b*. The PLAZA protein database was searched with the BLASTP method using the amino acid sequences of the Arabidopsis BB (AT3G63530). Two maize *BB* homologues were found in maize: GRMZM2G141084 for *BB1* and GRMZM2G021498 for *BB2* (Fig. S1B). Similar to the Arabidopsis BB protein, both maize BB proteins harbor the RING domain, and the protein sequences of these two maize BB showed a very high similarity to each other (90.4%) (Fig. S1C) suggesting that they might be functionally redundant.

### 3.2 Generation and phenotypic analysis of *DA1^R333K^* dominant-negative overexpression lines

Compared with the Arabidopsis DA1, the maize DA1 also contains a LIM domain, two UIM domains, a conserved C-terminal region, and the conserved arginine at 333^rd^ amino acid (Fig. S2). To characterize the function of maize *DA1*, we ectopically expressed the B104 *DA1^R333K^* under the control of the maize ubiquitin promoter into the maize inbred line B104. It was expected that this mutant *DA1* allele would interfere in a dominant-negative fashion with DA1 and DA1-related protein activity as it does in Arabidopsis [10]. Nine independent lines were obtained and backcrossed to B104 to obtain the BC_1_ segregating plants (WT and *DA1^R333K^* heterozygote). The BC_1_ shoots were used to select plants with a single-locus insertion based on the 1:1 Mendelian segregation ratio of the transgene, and five lines were found to contain a single locus insertion. To examine the expression level of the transgene in these five lines, total RNA was extracted from the first centimeter at the basis of the 4^th^ leaf, 2 days after its appearance, when the leaf was actively growing from transgenic plants and their counterpart segregating WT. The total transcripts of endogenous *DA1* and transgenic *DA1^R333K^* were significantly increased in these transgenic lines. *DA1^R333K^-3, −7* and −*9*, which showed the highest *DA1* expression among these transgenic lines were selected for further phenotypic analysis (Fig. S3).

For maize leaf growth, two key leaf growth parameters are important: the leaf elongation rate or maximal growth rate and the leaf elongation duration or how long maximal growth is maintained. These parameters can be determined by daily measuring the leaf length until it is maximal [31]. The final 4^th^ leaf length and LER of *DA1^R333K^-3* and −*9* were not significantly changed while these parameters were significantly decreased in *DA1^R333K^-7* (Fig. 1). Compared with WT, the final 4^th^ leaf length and LER of *DA1^R333K^-7* were reduced by 7.3% and 8.2%, respectively. Of the three lines with ectopic expression of *DA1^R333K^*, line 7 had the lowest expression levels (Fig. S3). The LED of three overexpression lines did not show a significant change (Fig. 1). Next, we measured some agronomic traits of the mature plants. The final plant height, ear leaf length and width, stem width and total leaf number of the overexpression plants were all similar to those of their segregating WT (Table 1).

**Figure 1.**
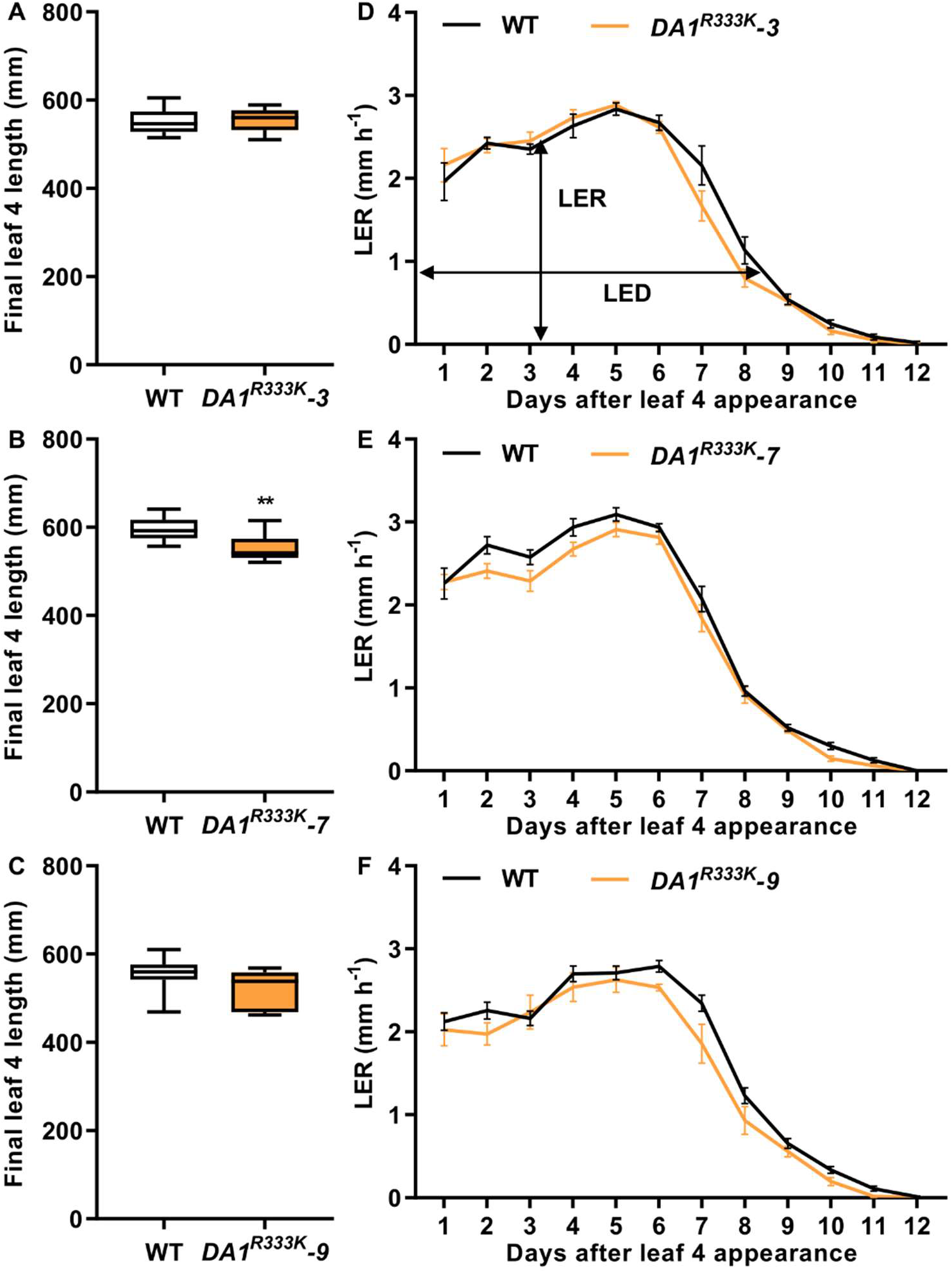
The final 4^th^ leaf length and LER curve of the *DA1^R333K^* overexpression lines. The final 4^th^ leaf length of *DA1^R333K^-3* (**A**), *DA1^R333K^-7* (**B**), and *DA1^R333K^-9* (**C**). The LER of *DA1^R333K^-3* (**D**), *DA1^R333K^-7* (**E**), and *DA1^R333K^-9* (**F**). In the box plots, the boundary of the box closest to zero indicates the 25^th^ percentile, a black line within the box marks the mean, and the boundary of the box farthest from zero indicates the 75^th^ percentile. Whiskers above and below the box indicate the max and min value. Error bars in the line charts represent standard error. Significant differences were determined using Student’s t-test: *, P<0.05, **, P<0.01, n≥8.

**Table 1.**
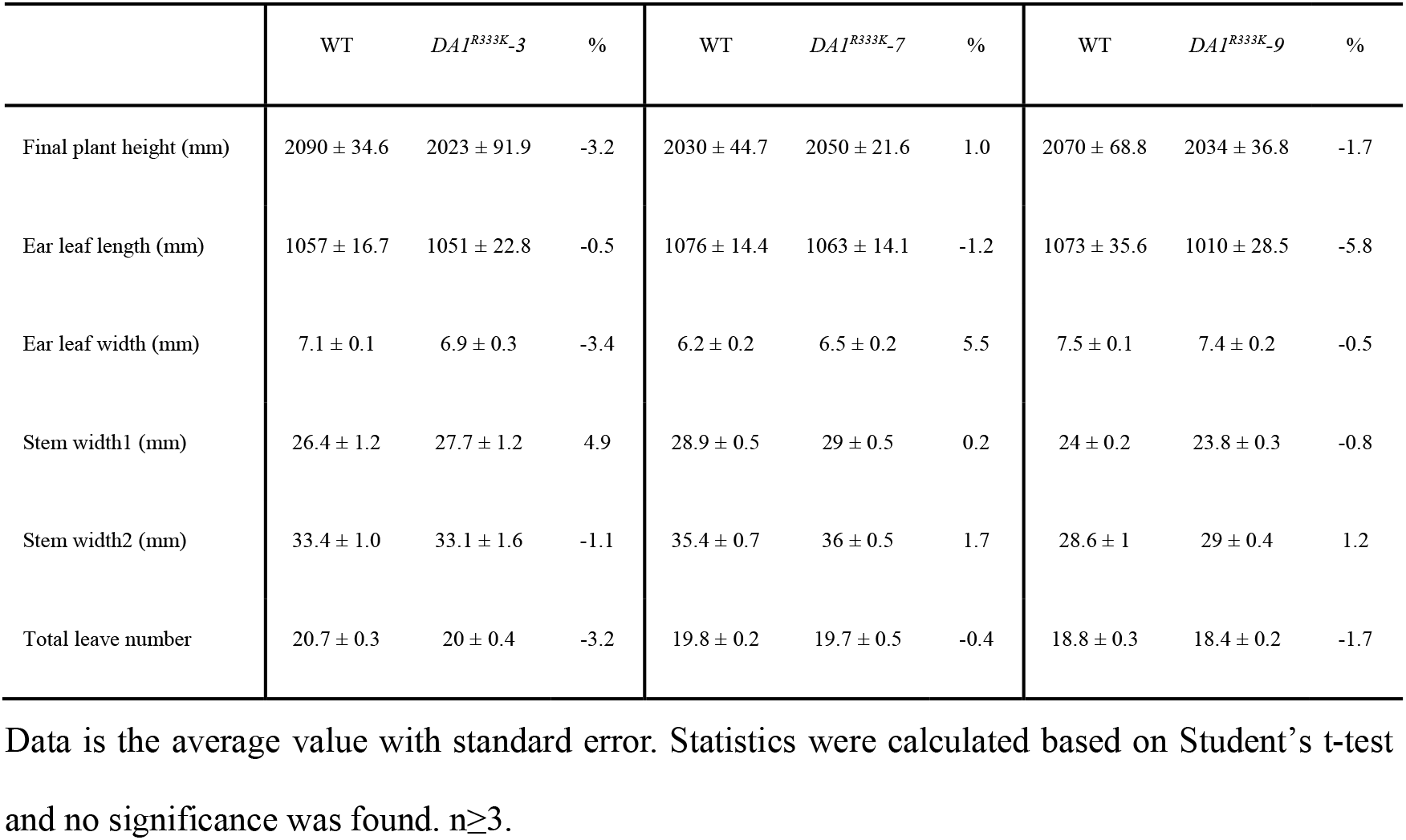
Overview of the phenotypes of WT and *DA1^R333K^* overexpression lines.

|  | WT | *DAI<sup>R333K</sup>-3* | % | WT | *DAI<sup>R333K</sup>-7* | % | WT | *DAI<sup>R333K</sup>-9* | % |
| --- | --- | --- | --- | --- | --- | --- | --- | --- | --- |
| Final plant height (mm) | 2090 ± 34.6 | 2023 ± 91.9 | -3.2 | 2030 ± 44.7 | 2050 ± 21.6 | 1.0 | 2070 ± 68.8 | 2034 ± 36.8 | -1.7 |
| Ear leaf length (mm) | 1057 ± 16.7 | 1051 ± 22.8 | -0.5 | 1076 ± 14.4 | 1063 ± 14.1 | -1.2 | 1073 ± 35.6 | 1010 ± 28.5 | -5.8 |
| Ear leaf width (mm) | 7.1 ± 0.1 | 6.9 ± 0.3 | -3.4 | 6.2 ± 0.2 | 6.5 ± 0.2 | 5.5 | 7.5 ± 0.1 | 7.4 ± 0.2 | -0.5 |
| Stem width1 (mm) | 26.4 ± 1.2 | 27.7 ± 1.2 | 4.9 | 28.9 ± 0.5 | 29 ± 0.5 | 0.2 | 24 ± 0.2 | 23.8 ± 0.3 | -0.8 |
| Stem width2 (mm) | 33.4 ± 1.0 | 33.1 ± 1.6 | -1.1 | 35.4 ± 0.7 | 36 ± 0.5 | 1.7 | 28.6 ± 1 | 29 ± 0.4 | 1.2 |
| Total leaf number | 20.7 ± 0.3 | 20 ± 0.4 | -3.2 | 19.8 ± 0.2 | 19.7 ± 0.5 | -0.4 | 18.8 ± 0.3 | 18.4 ± 0.2 | -1.7 |

Data is the average value with standard error. Statistics were calculated based on Student's t-test and no significance was found. n≥3.

To obtain homozygote plants, the BC_1_ heterozygotes were self-pollinated, and the BC_1_F_1_ WT and homozygous transgenic BC_1_F_1_ plants were self-pollinated to obtain the BC_1_F_2_ generation. The *DA1^R333K^-9* line which shows the highest expression of *DA1* was chosen for further analysis. The *DA1* expression level was increased by 56 times in the leaf basis of the homozygous *DA1^R333K^-9* shoots compared with WT (Fig. S4A). To investigate whether the homozygous *DA1^R333K^-9* plants affected leaf growth, a growth analysis of the 4^th^ leaf was conducted of plants grown under well-watered conditions (WW) as well as under water-deficient conditions (WD), by with-holding water the moment leaf 3 appeared. The results showed that under WW conditions, the homozygous *DA1^R333K^-9* shoots showed similar final 4^th^ leaf length and LER to WT (Fig. S4B-C). Furthermore, both WT and homozygous *DA1^R333K^-9* showed a similar and significant reduction of 4^th^ leaf length and LER under water-deficient conditions compared with WW conditions, demonstrating that the drought treatment had the intended negative effect on 4^th^ leaf growth. In conclusion, despite the high conservation of the arginine at position 333 and the high ectopic expression of *DA1^R333K^*, no reproducible effects on plant growth parameters could be observed, indicating that the DA1 pathway has a less prominent role in growth control than is the case for Arabidopsis.

### 3.3 Generation and phenotypic analysis of *da1_dar1a_dar1b* triple CRISPR mutants

In Arabidopsis, the single *da1* or *dar1* knockout mutants did not or slightly promote plant organ growth while the *da1_dar1* double knockout mutants showed larger flowers and seeds [10, 12]. In maize, there were two *DAR1* homologues that are closest related to *DA1* (Fig. S1A), sharing 68% similarity to each other. To further decipher the function of the maize *DA1* and *DAR1* genes, we generated *da1_dar1a_dar1b* knockout maize mutants by using CRISPR/Cas9 with a dual gRNA approach. The first gRNA targets the first exon of *DA1* (located on chromosome 1) and the second gRNA targets both *DAR1* genes (both located on chromosome 6) (Fig. 2A). Five independent transgenic lines were obtained including two lines contained mutations in all three genes, namely *da1_dar1a_dar1b-a* and *da1_dar1a_dar1b-c*. *DA1* showed, at target 1 site, a two bases deletion in the *da1_dar1a_dar1b-a* line and has a 62 bases deletion in the *da1_dar1a_dar1b-c* line, respectively. *DAR1a* had the same 4 bases deletion at target 2 site in both lines. *DAR1b* showed 4 bases deletion at target 2 site in both lines but at a different position (Fig. 2B). The frameshift mutations in the *DA1*, *DAR1a* and *DAR1b* caused premature stop codons that appeared before the first UIM domain in *da1* and before the conserved C-terminal region in *dar1a* and *dar1b*. The T_0_ mutants were backcrossed with B104 and then self-pollinated twice to obtain the BC_1_F_2_ generation for phenotypic analysis.

**Figure 2.**
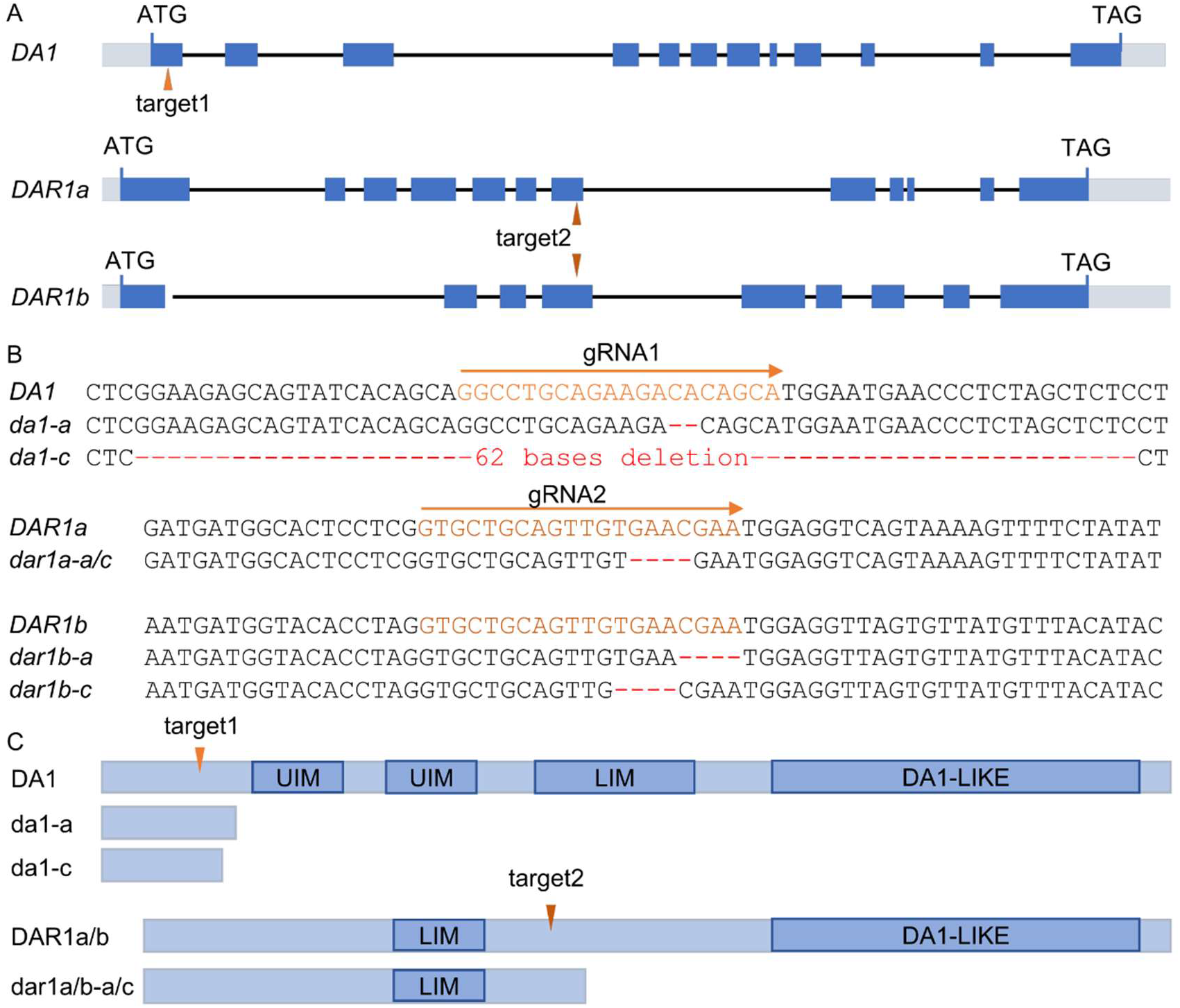
*da1_dar1a_dar1b* mutants obtained through CRISPR/Cas9 gene editing. **(A)** Structural representation of the maize *DA1, DAR1a* and *DAR1b* genes showing the target sites of the two gRNAs. **(B)** The gRNA sequences (orange) and mutation sites of the mutated alleles. **(C)** The protein structures of DA1, DAR1a and DAR1b and their putative mutant isoforms. The dark blue boxes represent the protein domains. “/” represents “or”.

At the seedling stage, there were little differences in final 4^th^ leaf length (−0.8%) and LER (1.4%) between *da1_dar1a_dar1b-a* triple mutants and WT (Fig. 3A-B). The mature *da1_dar1a_dar1b-a* triple mutant showed, compared with WT, a similar total leaf number, a slightly increased ear leaf length (2.4%) and stem width (9.8% and 12.2% depending on the direction of the elliptic internode). Final plant height of *da1_dar1a_dar1b-a* was significantly decreased by 7.7% (Table 2, Fig. 3C), while final plant height of *da1_dar1a_dar1b-c* plants was only slightly and not-significantly decreased by 0.3%. Conversely, the ear leaf length of *da1_dar1a_dar1b-c* was significantly increased by 4.6% (Table 2). For both triple mutants, the cob length and width were similar to WT and the seed area was slightly decreased in the triple mutant (−14.6% in *da1_dar1a_dar1b-a* and −2.0% in *da1_dar1a_dar1b-c*) (Fig. 3D, Table 2).

**Figure 3.**
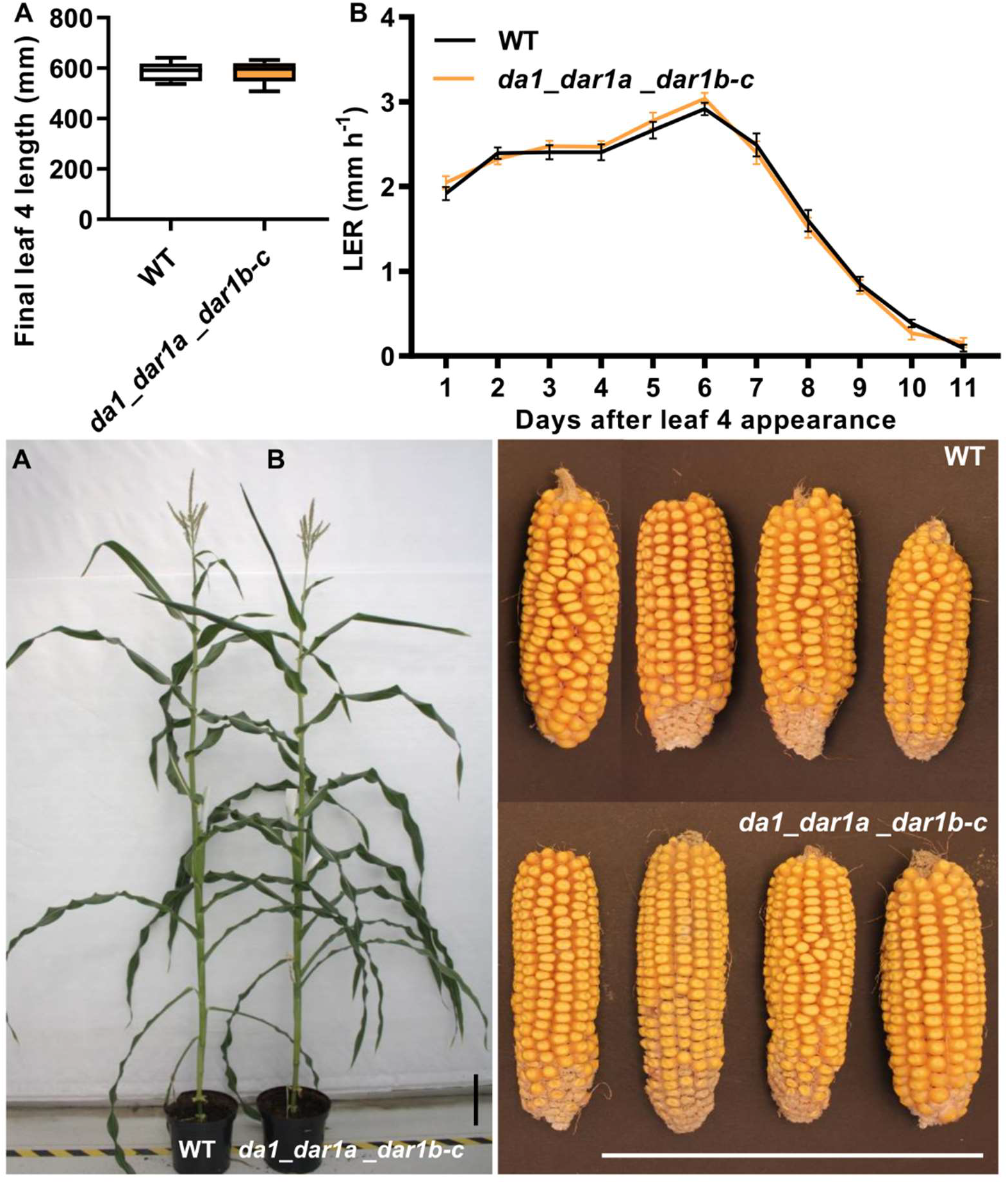
Overview of the phenotypes of WT and *da1_dar1a_dar1b* triple mutant plants. The final 4^th^ leaf length **(A)**, LER **(B)**, mature plants **(C)**, and cobs **(D)** of WT plant and triple mutants. In the box plots, the boundary of the box closest to zero indicates the 25^th^ percentile, a black line within the box marks the mean, and the boundary of the box farthest from zero indicates the 75^th^ percentile. Whiskers above and below the box indicate the max and min value. Error bars in the line charts represent standard error. Significant differences were determined using Student’s t-test: *, P<0.05, **, P<0.01, n≥8. Scale bar = 20 cm.

**Table 2.** Overview of the phenotypes of WT and *da1_dar1a_dar1b* triple mutants.

|  | WT | *dal_dar1a_dar1b-a* | % | P-value | WT | *dal_dar1a_dar1b-c* | % | P-value |
| --- | --- | --- | --- | --- | --- | --- | --- | --- |
| Final plant height (mm) | 2186.3 ± 50.7 | 2018.75 ± 23.3 | -7.7 | 0.04 | 2315.2 ± 74.1 | 2308 ± 39.7 | -0.3 | 0.94 |
| Ear leaf length (mm) | 1060.8 ± 12.3 | 1086.3 ± 10.1 | 2.4 | 0.19 | 1020.3 ± 14.7 | 1066.8 ± 10.1 | 4.6 | 0.04 |
| Ear leaf width (mm) | 73 ± 3.3 | 73 ± 1.9 | 0.0 | 1 | 72 ± 2.5 | 72 ± 201 | 0.0 | 1.00 |
| Stem width1 (mm) | 15.2 ± 0.7 | 16.7 ± 0.7 | 9.80 | 0.16 | 16.3 ± 0.7 | 16.9 ± 0.5 | 4.0 | 0.37 |
| Stem width2 (mm) | 15.5 ± 0.6 | 17.4 ± 0.8 | 12.2 | 0.1 | 17.6 ± 0.7 | 16.2 ± 0.6 | -7.5 | 0.39 |
| Total leaf number | 21.8 ± 0.25 | 21 ± 0.4 | -3.5 | 0.18 | 22.5 ± 0.5 | 21.6 ± 0.5 | -4.0 | 0.25 |
| Seed area (mm2) | 55 ± 3.6 | 47 ± 3.9 | -14.6 | 0.14 | 49 ± 0.5 | 48 ± 2.3 | -2.0 | 0.56 |

Data is the average value with standard error. Statistics were calculated based on Student's t-test. n≥4.

To analyze whether the triple mutants are tolerant to drought stress, *da1_dar1a_dar1b-c* seedlings and WTs were subjected to WW and a mild drought (MD) treatment (water content of the soil is 70% of the WW conditions). The results showed that the 4^th^ leaf length and shoot fresh weight of the triple *da1_dar1a_dar1b-c* mutants were decreased by 0.2% and 10.6% (albeit not significant) respectively, compared with WT under WW conditions (Fig. 4 A-B). Under MD conditions, the 4^th^ leaf length of *da1_dar1a_dar1b-c* was slightly increased by 0.5% while the shoot fresh weight was decreased by 15.4% (both not significant). Compared with WW, the 4^th^ leaf length of WT and *da1_dar1a_dar1b-c* mutants was significantly decreased under MD conditions due to a decreased LER (Fig. 4C). The 4^th^ leaf length of WT was reduced by 10.0% in MD, while the *da1_dar1a_dar1b-c* mutants decreased by 9.3%. The shoot fresh weight of WT decreased by 63.8% in response to the drought treatment, very comparable to the 65.8% reduction observed in the triple mutants (Fig. 4A-B). Altogether, these results suggested that plant development of the *da1_dar1a_dar1b* triple mutants was only marginally affected in the conditions analyzed, with a possible slight negative effect on plant height. There is no significant effect of inactivation of the *DA1*, *DAR1a* and *DAR1b* on seedling drought tolerance.

**Figure 4.**
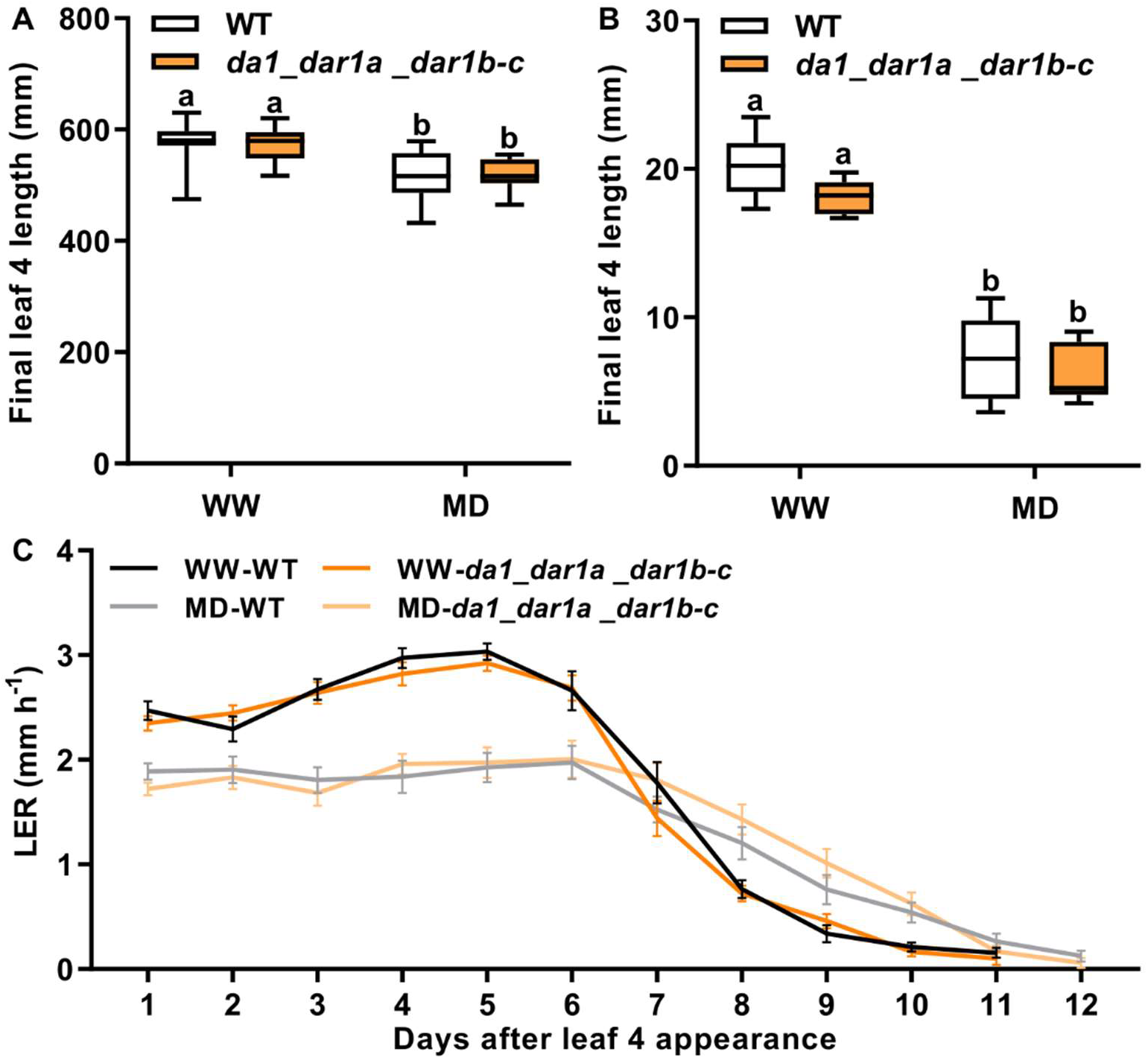
Leaf four growth phenotype of WT and *da1_dar1a_dar1b* triple mutant under mild drought conditions. The final 4^th^ leaf length **(A)**, shoot fresh weight **(B)**, and LER **(C)** of WT plant and triple mutants. In the box plots, the boundary of the box closest to zero indicates the 25^th^ percentile, a black line within the box marks the mean, and the boundary of the box farthest from zero indicates the 75^th^ percentile. Whiskers above and below the box indicate the max and min value. Error bars in the line charts represent standard error. Different letters indicate P < 0.05 according to Tukey’s test (n≥8).

### 3.4 Generation and phenotypic analysis of *bb1_bb2* double CRISPR mutants

Because of the high similarity of the two *BB* genes, we designed two gRNAs that target both genes, one at the 5^th^ exon and another at the 7^th^ exons (Fig. 5A). The h-line had a large deletion (including an intron) between 2 gRNAs in both genes. The m-line had a biallelic mutation at target 1 (one base and two bases insertion) and one base insertion at target 2 in *BB1* and contains one base insertion at both targets in *BB2* (Fig. 5B). Due to the large deletion in the h-line and the frameshift caused by one or two bases insertions in m-line, the *bb1* and *bb2* mutants lost the entire RING domain, which is essential for interacting with the E2 (Fig. 5C).

**Figure 5.**
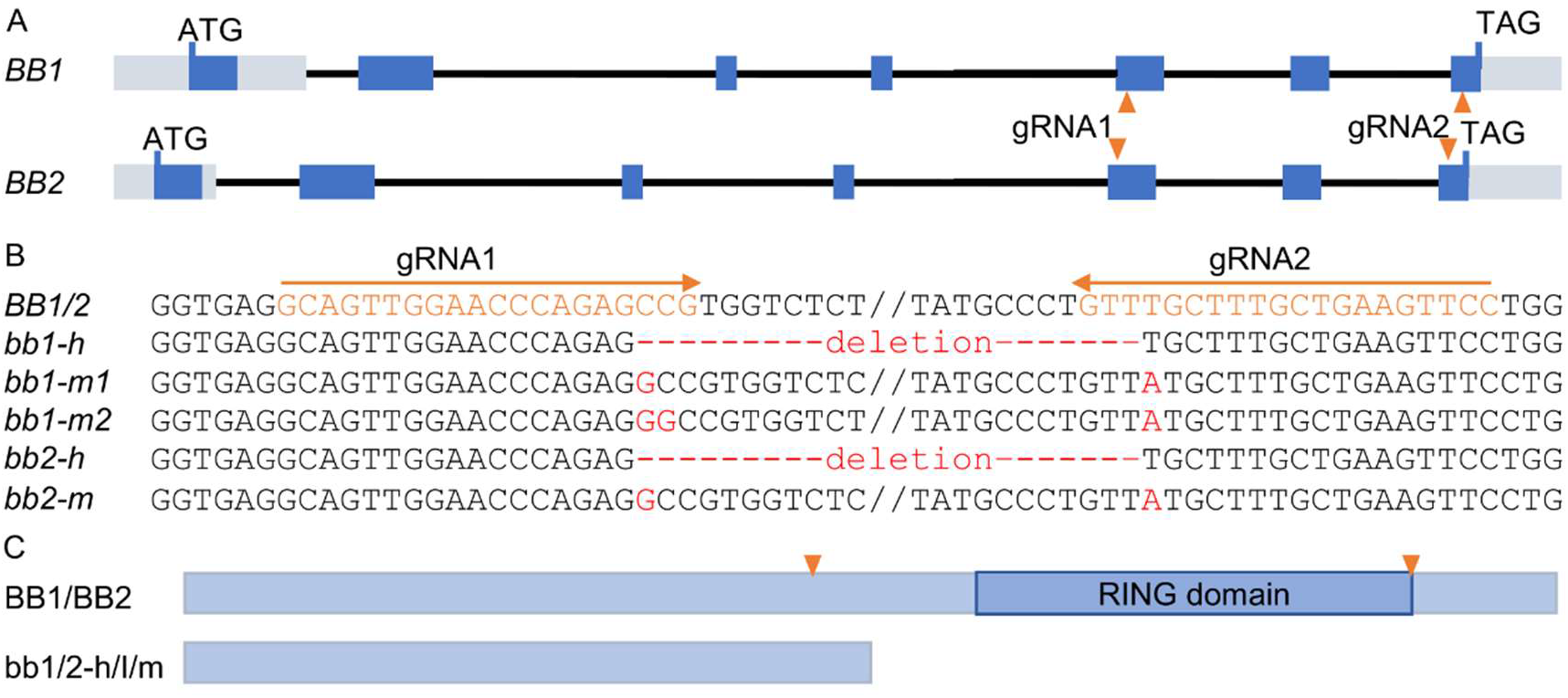
bb1 and bb2 mutants obtained through CRISPR/Cas9 gene editing. **(A)** Structural representation of the maize *BB1* and *BB2* genes showing the target sites of the two gRNAs. **(B)** The gRNA sequences (orange) and mutation sites of the mutated alleles. **(C)** The protein structures of the BB genes and their putative mutant isoforms. The dark blue boxes represent the protein domains. “/” represents “or”.

BC_1_F_2_ plants of the h-line and m-line (the allele contains two bases insertion at target 1 of BB1) were used for phenotyping. The final 4^th^ leaf length and LER of *bb1, bb2*, and *bb1_bb2* double mutants of both lines were not significantly affected compared with WT (Fig. 6A-D). Agronomic traits such as plant height, ear leaf size, stem width and leaf number also were in the same range as those of WT (Fig. 6E and Table 3). We further analyzed the seed weight and seed size. For the h-line, 100 seeds weight of the *bb1, bb2* and *bb1_bb-2* mutants was slightly increased by 8.5%, 8.5% and 9.6%, respectively, compared with WT. Also, the seeds area of *bb1, bb2* and *bb1_bb-2* mutants were enlarged by 13.4%, 10.8% and 12.2%, respectively. However, these increases were not statistically significant due to a large individual variation within the genotypes. For the m-line, the 100 seed weight of *bb1, bb2* and *bb1_bb-2* double mutants were altered by 11.9%, −1.0%, and 0%, respectively; while the seed area of *bb1, bb2* and *bb1_bb-2* mutants was reduced by 0.1%, 7.3%, and 5.4% (Table 3). In addition, the double mutant of the h-line was exposed to drought conditions. Both WT and double mutant in WD conditions showed a significantly reduced 4^th^ leaf length compared to their counterparts in WW conditions, the reduction of WT is 13.7% while and it is 10.9% in the double mutant (Fig.7). However, the inactivation of both BB1 and BB2 did not significantly alter the 4^th^ leaf length, LER or LED compared with WT plants when grown under the same conditions.

**Figure 6.**
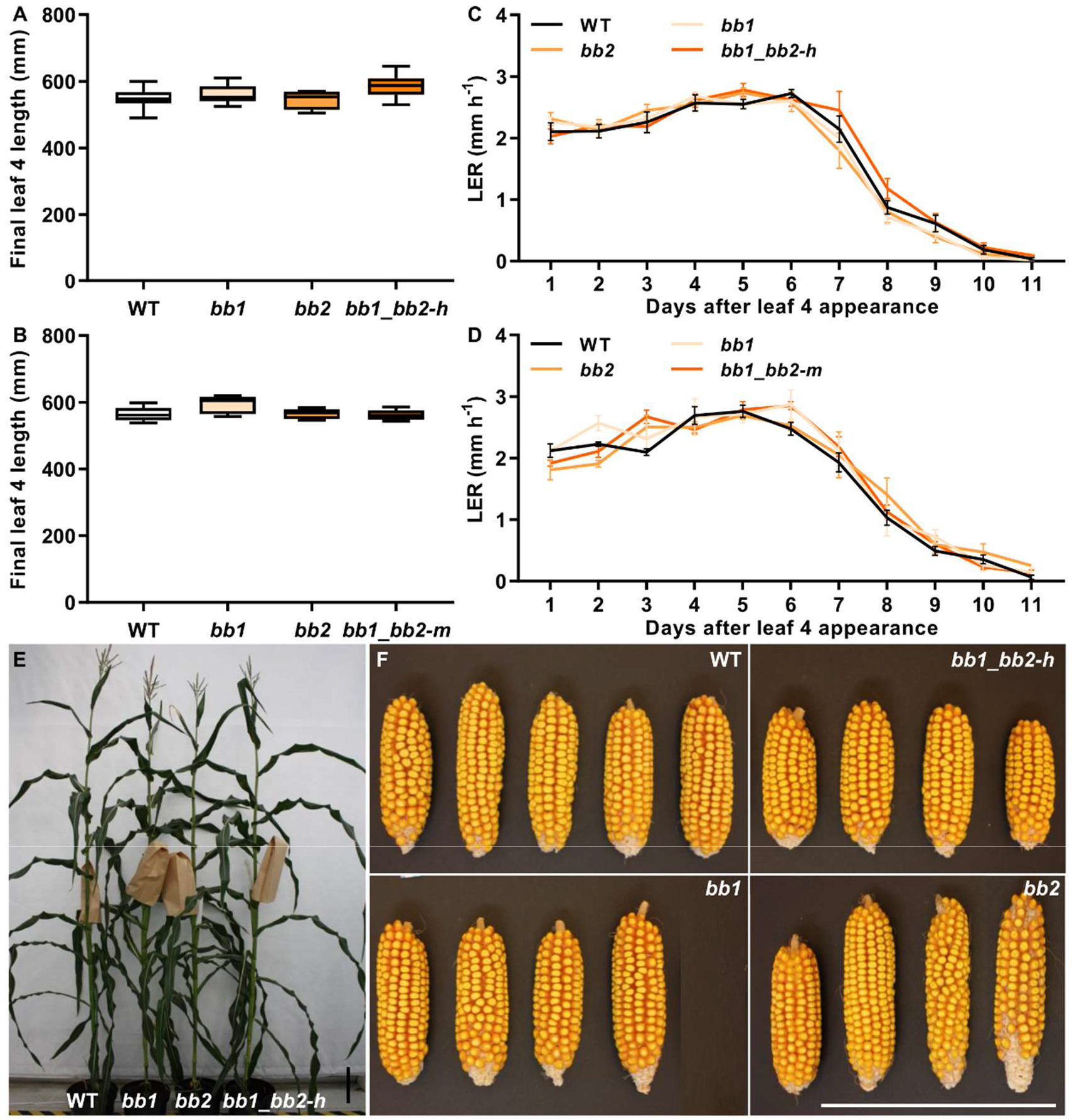
Overview of the phenotypes of WT and *bb1_bb2* double mutant plants. The final 4^th^ leaf length of h line**(A)**, of m line **(B)** the LER of h line **(C)**, of m line **(D).** The mature plants **(E)**, and cobs **(F)** of WT plant and *bb* mutants. In the box plots, the boundary of the box closest to zero indicates the 25^th^ percentile, a black line within the box marks the mean, and the boundary of the box farthest from zero indicates the 75^th^ percentile. Whiskers above and below the box indicate the max and min value. Error bars in the line charts represent standard error. No significant differences were found according to Tukey’s test, n≥8. Scale bar = 20 cm.

**Figure 7.**
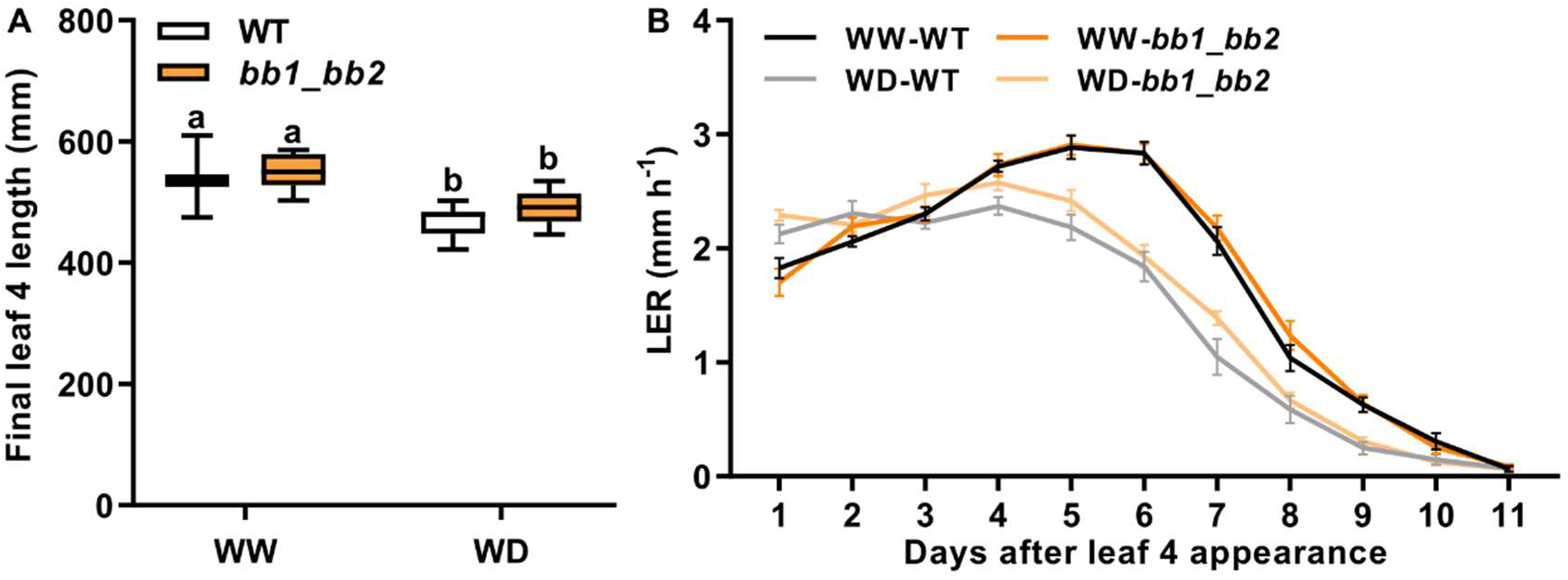
Leaf four growth phenotype of WT and *bb1_bb2* mutant under drought conditions. The final 4^th^ leaf length **(A)** and LER **(B)** of WT plant and *bb1_bb2* mutant. In the box plots, the boundary of the box closest to zero indicates the 25^th^ percentile, a black line within the box marks the mean, and the boundary of the box farthest from zero indicates the 75^th^ percentile. Whiskers above and below the box indicate the max and min value. Error bars in the line charts represent standard error. Different letters indicate P < 0.05 according to Tukey’s test (n≥8).

**Table 3.**
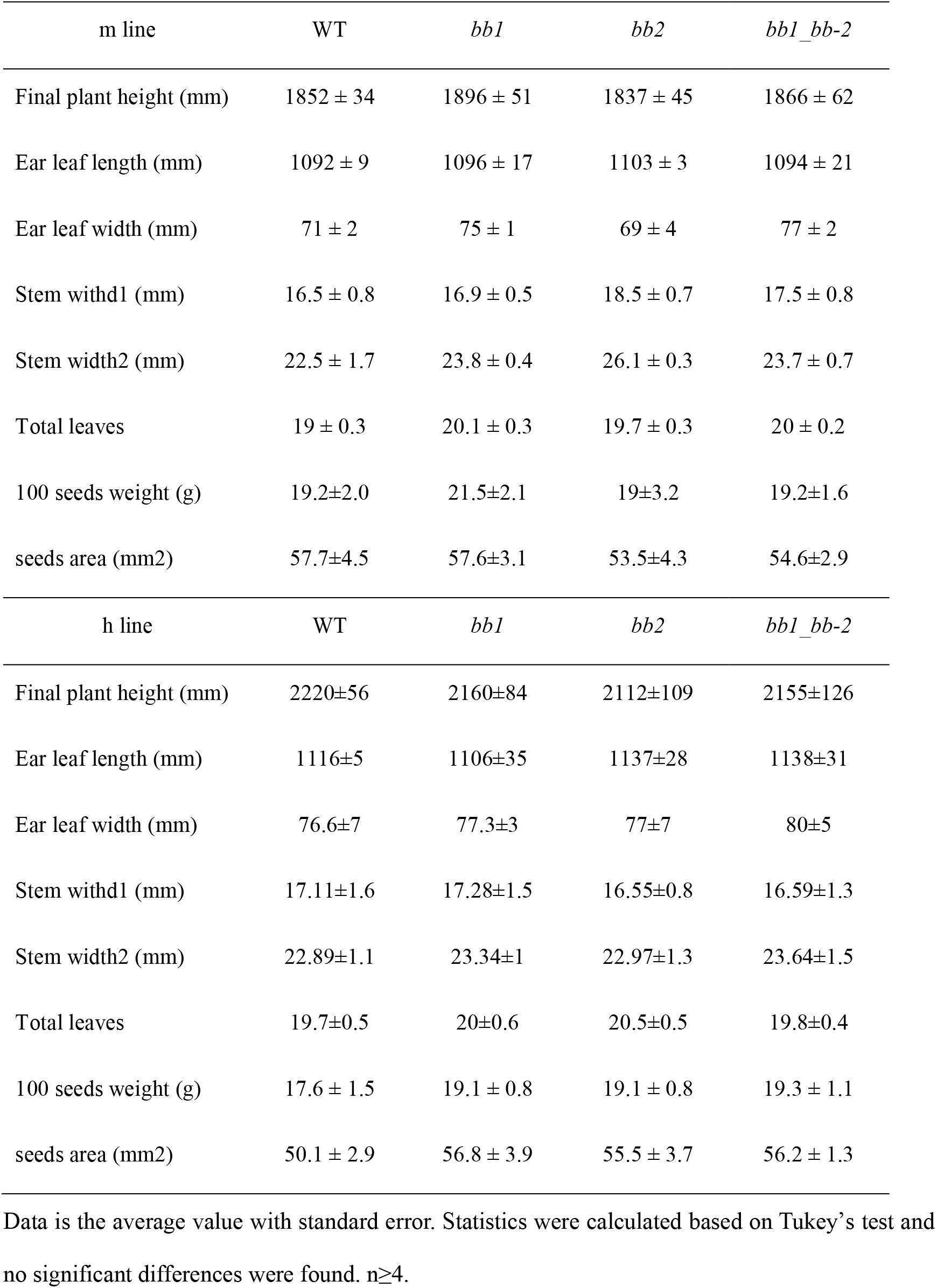
Overview of the phenotypes of WT and bb mutants.

In Arabidopsis, the dominant *da1-1* mutation is known to enhance the phenotype of *bb* mutant [10, 12, 13]. Therefore, we also crossed the maize *DA1^R333K^-9* with the *bb1_bb-2* mutant (h-line) to generate the *DA1^R333K^_bb1_bb2* plants. Compared with WT, the *DA1* expression in the plants was still significantly increased (Fig. S5). The final plant height of the *DA1^R333K^_bb1_bb2* was significantly decreased (10.8%) compared with WT. The 4^th^ leaf length, ear leaf size, stem width and leaf number of the *DA1^R333K^_bb1_bb2* were similar to WT (Fig. 8, Table 4). Altogether, these results indicated that modification of the *DA1, DAR1* and *BB* genes in maize did not obviously promote organ size and drought tolerance of seedlings.

**Figure 8.**
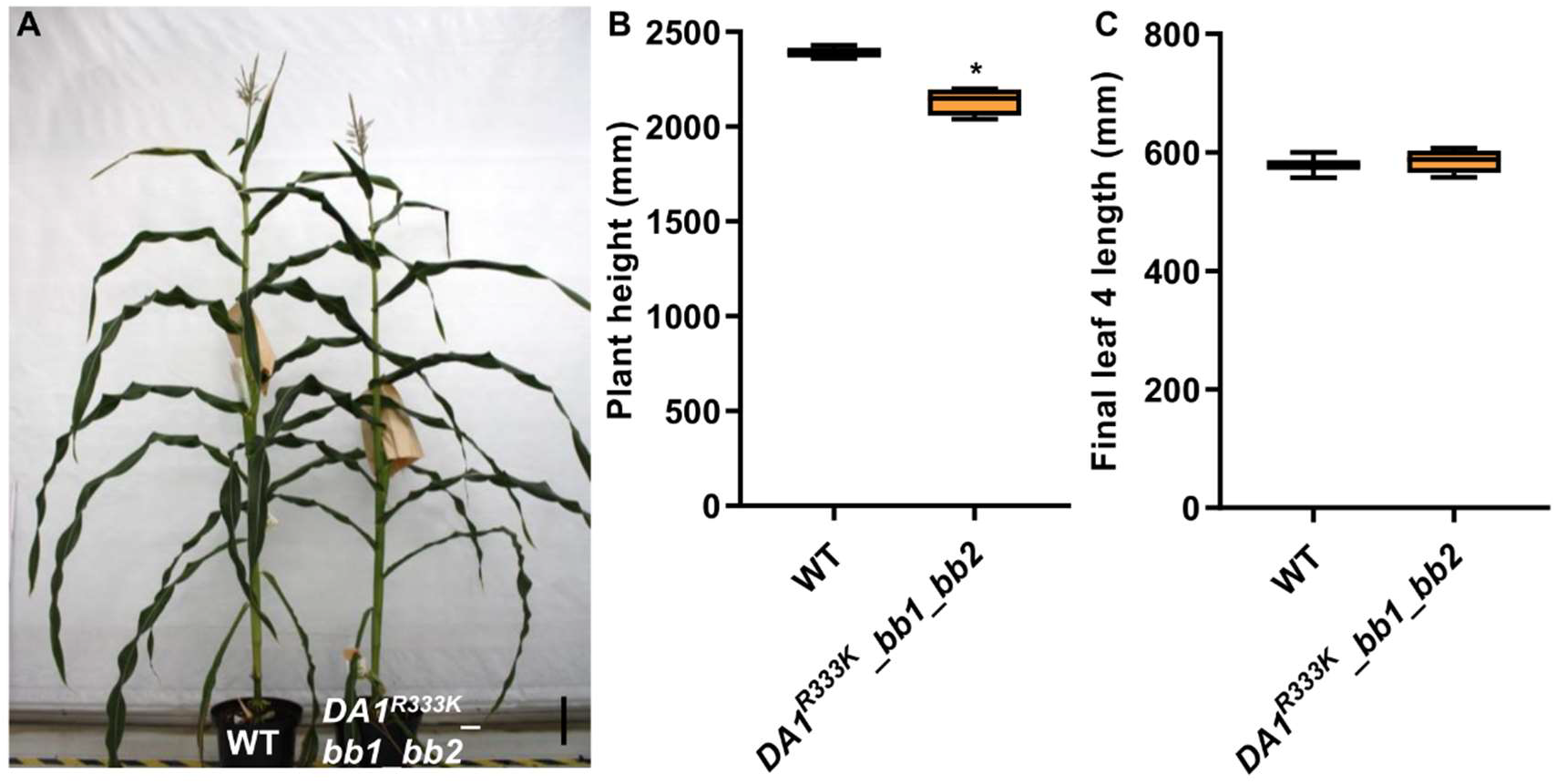
Overview of the phenotypes of WT and *DA1^R333K^ _bb1_bb2* mutant plants. The mature plants **(A)**, plant height **(B)** and 4^th^ leaf length **(C)** of WT plant and *DA1^R333K^ _bb1_bb2* mutants. In the box plots, the boundary of the box closest to zero indicates the 25^th^ percentile, a black line within the box marks the mean, and the boundary of the box farthest from zero indicates the 75^th^ percentile. Whiskers above and below the box indicate the max and min value. Significant differences were determined using Student’s t-test: *, P<0.05, n≥4. Scale bar = 20 cm.

**Table 4.** Overview of the phenotypes of WT and *DA1^R333K^_bb1_bb2* mutants.

|  | WT | *DAI<sup>R333K</sup>_bb1_bb2* | % | P-value |
| --- | --- | --- | --- | --- |
| Final plant height (mm) | 2393.3 ± 20.3 | 2135 ± 35.9 | -10.8 | 0.002 |
| Ear leaf length (mm) | 1032.5 ± 15.5 | 1036.5 ± 55.8 | 0.4 | 0.93 |
| Ear leaf width (mm) | 76.2 ± 1.26 | 75 ± 4.69 | -1.6 | 0.8 |
| Stem width1 (mm) | 15.2 ± 0.7 | 14.3 ± 0.6 | -5.8 | 0.33 |
| Stem width2 (mm) | 15.5 ± 0.4 | 15 ± 1 | -2.9 | 0.6 |
| Total leaves | 21 ± 0 | 20.8 ± 0.5 | -1.2 | 0.39 |

Data is the average value with standard error. Statistics were calculated based on Student's t-test. n≥3.

## 4. Discussion

In Arabidopsis, seven DARs, denominated AtDAR1 to AtDAR7, show similarity to the AtDA1 protein sequence, of which at least DA1, AtDAR1, and AtDAR2 are functionally redundant [10, 14]. In maize, homologues of *DA1* and only five *DARs* were found and all of them encoding proteins harbor the LIM domain and the conserved C-terminal region [18]. Two of the maize *DAR* genes cluster together with the *AtDAR1* and another two *DAR* genes were close to the *AtDAR2* (Fig. S1A). The *AtDA1, AtDAR1* and *AtDAR2* showed a similar expression pattern with the strongest expression in the developing young leaves [10, 14]. In maize, the *DA1* and *DARs* were ubiquitously expressed during maize development. In contrast to the Arabidopsis *AtDA1*, *AtDAR1*, and *AtDAR2*, the maize *DA1, DAR1s* and *DAR2s* are more highly expressed in mature leaves as compared to developing leaves (Table S1). These data indicated that these genes may also participate in later stages of leaf development.

Whereas the role of *DA1* and *DAR* genes in leaf development is clear in Arabidopsis, it is far from obvious in maize. Neither transgenic maize plants that express a dominant-negative *DA1^R333K^* allele nor a maize *da1_dar1a_dar1b* triple mutant showed a robustly altered organ or leaf size. Possibly, the two maize *DAR2* genes are sufficient to sustain growth in the *DA1^R333K^* overexpression line and the *da1_dar1a_dar1b* triple mutant. To address this hypothesis, a pentuple *da1_dar1a_dar1b_dar2-1_dar2-2* mutant needs to be made and phenotypically analyzed. Moreover, ectopic overexpression of the DA1^R358K^ dominant-negative allele in 17 diverse Arabidopsis accessions caused a strong to moderate increased leaf size in the different accessions indicating that the phenotype of *DA1^R358K^* is influenced by genetic background [13]. Our data are consistent with the observation that overexpression of dominant-negative alleles of *DA1* or *DAR1* in the maize inbred line DH4866 did not promote final leaf size [18]. However, in the latter study, an increase in seed size was observed, a phenotype that was not noticed in the current study. A possible explanation could be that the effects of DA1 and DAR proteins on regulating organ size depend on the genetic background of the maize line and that the phenotypes are less penetrant in B104 that was used in this study.

*BB* and *DA2* are two well-characterized *DA1* regulators in Arabidopsis and both genes were highly expressed in young tissues [19, 20]. Mutations in *BB* or *DA2* showed enlarged organ size due to an increased cell number [19, 20]. In Arabidopsis, BB and DA2 physically interact with DA1 and regulating the activity of DA1 and DAR1 through ubiquitination [10, 19]. Both *da1-1_bb* and *da1-1_da2* double mutants displayed synergistic effects on the size of seed and petal, while the *bb_da2* double mutants showed an additive effect on organ size, suggesting that the two E3 ligases worked independently [10, 12, 13, 19]. Here, we found two maize *BB* homologues, and both *BB* genes were highly expressed in the young proliferating tissues such as SAM, immature leaf, and seeds, suggesting the maize *BB* genes may also be involved in regulating cell proliferation (Table S1). To study the role of these *BB* genes in organ development and size control, we generated *bb1_bb2* double mutants by using CRISPR/Cas9. However, phenotypic analysis of two independent maize lines in which both BB genes were inactivated did not show a significantly increased leaf, stem or seed size compared with WT. Maybe, in maize and in contrast to Arabidopsis, the functions of *BB* and *DA2* are functionally redundant. Moreover, the plant height of *DA1^R333K^ bb1_bb2* plants were significantly decreased while no significant changes in the size of leaves, indicating that the *DA1* and *BB* genes may also involve in regulating maize organ size and the effects might be organ-specific. This organ-specific effects of *DA1* genes were also observed in Arabidopsis. For instance, the Arabidopsis triple *da1-ko1_dar1-1_dar2-1* mutant showed enlarged seeds and petals, while the size of leaves were extremely decreased [14]. Furthermore, our work is a clear demonstration of the difficulties of translating observations made in model plants, such as Arabidopsis, to crops, such as maize.

Most maize traits are controlled by many small-effect genes [32–35], and to obtain desirable traits may need to modify several genes. One of the advantages of CRISPR/Cas9 technology is multiple genes can be engineered simultaneously. Our study showed that CRISPR/Cas9 also works efficiently in editing multiple maize genes in single transformation experiments. Most of the mutations in *DA1, DAR1*, and *BB* genes were small insertions and deletions, also a large deletion between two gRNAs was observed in *BB* genes. Generating more mutants of *DA1* related genes such as *DAR2* and *DA2* using CRISPR/Cas9 would help us decipher the DA1 network in maize.

## Acknowledgements

The research was funded by the ‘Bijzonder Onderzoeksfonds Methusalem Project’ (no. BOFMET2015000201) of Ghent University. Pan Gong received a scholarship from the China Scholarship Council (CSC no. 201606320217).

---

**Figure S1. Phylogenic trees of DA1 families and BB genes. (B)** Phylogenic tree of *DA1* families in maize and Arabidopsis. **(B)** Phylogenic tree of *BB* and *BB* homologues in maize and Arabidopsis. **(C)** Amino acid sequences alignment of maize BB1 and BB2, the RING domain was marked with a box.

**Figure S2. The amino acid sequences alignment of Arabidopsis DA1 (AtDA1) and maize DA1 (ZmDA1) proteins.** The ubiquitin-interacting motif (UIM), LIM domain, DA1 peptidase motif and the mutated arginine were marked in the picture.

**Figure S3. The relative expression of DA1 in *DA1^R333K^* overexpression lines contains a single locus insertion.** Error bars represent standard error and significant differences were determined using Student’s t-test: *, P<0.05, **, P<0.01, n=3.

**Figure S4. Leaf four growth phenotype of WT and *DA1^R333K^-9* plants under drought conditions**. The relative expression of DA1 **(A)**, final 4^th^ leaf length **(B)** and LER **(C)** of WT plant and *DA1^R333K^-9* plants. Error bars in the line and bar charts represent standard error. Significant differences in the bar chart were determined using Student’s t-test: **, P<0.01, n=3. In the box plots, the boundary of the box closest to zero indicates the 25^th^ percentile, a black line within the box marks the mean, and the boundary of the box farthest from zero indicates the 75^th^ percentile. Whiskers above and below the box indicate the max and min value. Different letters in the box plot indicate P < 0.05 according to Tukey’s test (n≥6).

**Figure S5. The relative expression of DA1 in *DA1^R333K^_bb1_bb2* mutant**. Error bars represent standard error and significant differences were determined using Student’s t-test: *, P<0.05, **, P<0.01, n=3.

